# Destabilization of Structured RNAs by OPC and TIP4PD Water Models

**DOI:** 10.1101/2025.09.18.677163

**Authors:** Miroslav Krepl, Vojtěch Mlýnský, Pavel Banáš, Michal Otyepka, Jiří Šponer

**Author notes:** Corresponding author: Miroslav Krepl.

## Abstract

The four-point OPC water model has recently gained reputation as the preferred choice for molecular dynamics (MD) simulations of nucleic acids and even proteins, providing more realistic reproduction of bulk physical properties of water than the older three-point models. It has been shown to improve for example simulations of unstructured biomolecules such as RNA tetranucleotides or intrinsically disordered proteins. However, the performance for folded RNA structures was not specifically explored. In this study, we present extensive testing of the OPC water model on three different RNAs with highly intricate tertiary structures – the ribosomal L1 stalk RNA-protein protuberance, the mini tetraloop-tetraloop receptor (miniTTR-6) folded RNA, and the GAAA tetraloop-tetraloop receptor homodimer. The OPC performance is directly compared against SPC/E, TIP3P, OPC3, and TIP4P-EW water models along with the common OL3 AMBER RNA force field (FF). We found substantial effect of the water model on simulation behavior. For all three systems, we observe large-scale unfolding of the RNA, and even loss of the L1 stalk protein-RNA interface, when simulated with the OPC. In contrast, the simulations are entirely stable with the three-point water models. The underlying cause seems to be the higher affinity of the OPC waters to H-bond donor and acceptor groups of the RNA, disrupting the native solute-solute interactions. An identical issue can be observed also for the similar and widely used TIP4PD water model combined with the DES-Amber RNA FF. Our findings suggest that caution is warranted when using the four-point OPC and TIP4PD water models for simulations of structured RNAs. In combination with the current AMBER RNA FFs, the three-point water models may provide more realistic alternatives.

## Introduction

Due to their nature as a charged biopolymer, the nucleic acids can be sensitive to a water model chosen in explicit-solvent molecular dynamics (MD) simulations.^1-16^ An important part of the parametrization efforts aimed at improving the solute force-field (FF) performance for RNA have therefore been identifying the best performing water model.^2, 17-18^ The four-point OPC water model^19^ has recently come to be regarded as the optimal choice for nucleic acid MD simulations. When combined with RNA phosphate parameters refined by Steinbrecher et al.^20^ and the common OL3 AMBER RNA FF,^21^ the use of OPC improves description of single-stranded RNA molecules, such as tetranucleotides (TNs).^2^ This result has been confirmed in subsequent studies investigating RNA single-strands and tetraloops (TLs).^17-18, 22-27^. Recent improvements of the AMBER RNA FF and attempts to parameterize a new nucleic acid FF from the scratch are also made with the OPC water model.^22, 25-26, 28-29^ In addition, the AMBER ff19SB protein FF^30^ recommends the use of the OPC as well, due to its improved performance for intrinsically disordered proteins.^31^ However, beyond the TNs, TLs, and A-RNA duplexes, the OPC water model remains rather untested on more complex RNA systems, despite RNA’s ability to form highly intricate folded structures.^32^

In this work, we assess the performance of the four-point OPC water model on three different structured RNA systems and compare it with the other water models such as TIP3P,^33^ SPC/E,^34^ OPC3,^35^ and TIP4PEW.^36^ The common OL3 FF is used to describe the RNA in these simulations. We also evaluate another commonly used four-point water model, the TIP4PD,^37^ in combination with the DES-Amber RNA FF.^38^ Note that the design of four-point water models differs from that of three-point models by the presence of an additional off-atom charge site, intended to provide a more realistic electrostatic potential and, consequently, a more accurate representation of water bulk properties.

The first tested system, the ***L1 stalk***, is a flexible protuberance of the large ribosomal subunit, playing a crucial role in protein synthesis.^39^ It facilitates the release of deacylated tRNA after amino acid transfer and ensures smooth elongation cycle progression.^40-43^ Its apex rRNA structure (hereafter referred to as the L1 stalk rRNA) has a highly conserved and autonomously folding structure that is rich in non-canonical RNA interactions, including non-canonical base pairs and triplets, and base-phosphate, base-sugar and sugar-phosphate interactions.^44-45^ It also contains several recurrent RNA motifs, including two kink-turns^46^ and a TL.^47^ We have simulated the L1 stalk rRNA with as well as without the bound L1 protein.

The second tested system, the ***mini tetraloop–tetraloop receptor 6*** (miniTTR-6) structure, was specifically designed as a very stable structural element for RNA nanotechnology applications.^48^ While the miniTTR-6 lacks the biological significance of the ribosomal L1 stalk, the fact it has been deliberately engineered for high stability^48^ makes it a useful benchmark system as basically any reliable FF should maintain its structure over affordable timescales of standard MD simulations without major structural changes. The tetraloop-tetraloop receptor (TTR) interaction is the key long range tertiary contact in the system.^49^

The last of the tested systems is a simple symmetric homodimeric system consisting of two RNA duplexes interacting with each other via two equivalent TTR motifs.^50^ These two TTR motifs are the same one as in the miniTTR-6 and the system will be henceforth referred to as ***hTTR*** (homodimeric TTR).^51^ The hTTR was chosen as a testing system as its folding does not require the magnesium ions;^51^ in fact the isolated TTR forms in complete absence of magnesium merely in presence of moderate excess of KCl.^49^ There is an experimental evidence that even the miniTTR-6 system may not require the magnesium to fold in a sufficient excess of monovalent cations.^52^ All the studied systems are visualized in Supporting Information Figure S1.

Our standard simulations reveal that the OPC and TIP4PD water models destabilize key tertiary RNA-RNA interactions in all three systems compared to three-point water models. In many cases, this leads to visually striking structural changes of the RNA on unrealistically short timescales. For the L1 stalk, we even observe collapse of the protein-RNA interface that occurs independently of the RNA fold disruptions. The structural changes observed with the OL3/OPC combination are sometimes reversible on the simulation timescale whereas the DES-Amber/TIP4PD leads to more permanent disruptions. However, qualitatively identical structural changes are observed with both FF combinations. Based on detailed analysis of all systems and additional calculations directly comparing the OPC and TIP4PD water models to SPC/E, we propose the imbalance introduced by these four-point water models lies in their increased affinity to RNA H-bond donor and acceptor groups, disrupting the native RNA-RNA interactions. Though the difference is subtle for the individual H-bonds (probably ∼0.1–0.2 kcal/mol) and therefore likely negligible for small model systems, it may accumulate with the numerous interactions present in large structured RNAs and collectively determining the stability. For large folded RNA systems, this is apparently enough to significantly shift the free energy minima away from the native RNA fold towards more extended and therefore more solvated structures. In principle, all solute-solute H-bonds are affected by this. However the tertiary interactions formed by the RNA 2-OH′ groups are possibly the most sensitive and quickest to respond due to the complex balance between direct and water-mediated interactions. These interactions are omnipresent in large folded RNAs and include, e.g., the A-minor interaction^53^ – the most common tertiary RNA-RNA interaction in the ribosome. In conclusion, we suggest one of the three-point water models (e.g. TIP3P, SPC/E, OPC3) might be a more reliable choice for simulations of large structured RNAs than the currently popular four-point models.

## Methods

### Starting structures

We used the X-ray structure of the L1 stalk rRNA in complex with the L1 protein from *T. thermophilus* (PDB: 3U4M)^44^ as the starting structure for our simulations of the L1 stalk protein-RNA complex. Simulations of isolated L1 stalk rRNA were performed by removing the protein atoms. The cytidine 2111 was always N3-protonated in the starting structure. For simulations of the isolated L1 stalk rRNA, we also utilized its X-ray structure from *H. marismortui* (PDB: 5ML7).^54^ For the simulations of the miniTTR-6, we used its X-ray structure (PDB: 6DVK).^48^ Lastly, the first ensemble frame of the NMR structure of hTTR complex (PDB: 2I7Z) was used as the starting structure for its simulations.

All experimentally determined water molecules and monovalent ions were kept in the starting structures. Any experimentally detected monovalent ions other than K^+^ and Cl^-^ were converted into the latter ions. Experimentally determined (X-ray) divalent magnesium ions were utilized in selected simulations of the miniTTR6 system. For simulations where we included magnesium, any non-magnesium divalent cations were converted into magnesium prior to the system building; this resulted into nine bound Mg^2+^ ions in the starting structure of miniTTR6.

Besides simulations of folded RNAs, we have also performed simulations of the isolated cytidine nucleoside, with its structure taken from the AMBER residue library.

### Force-field selection

All coordinates and topology files were generated using the xLeap module of AMBER 22.^55^ The RNA systems were parameterized with the ff99bsc0χ_OL3_ (OL3) FF,^21, 56-57^ which is currently the recommended first-choice AMBER FF for RNA.^58^ In a few simulations, the OL3 FF was used with the adjusted phosphate van der Waals (vdW) parameters by Steinbrecher et al.;^20^ this version is abbreviated as OL3_CP_ (CP stands for “Case phosphate”).^20-21^ Note that in other literature and MD softwares, it is often referred to as “LJbb” FF. The reason for its testing was that the inclusion of adjusted phosphate vdW parameters together with the OPC water model was reported to somewhat improve the simulations of RNA TNs.^2^ However, the present simulations indicate that the CP modification does not have any effect on the instabilities observed for the folded RNAs tested here. For the protein component in the simulations of the L1 stalk protein-RNA complex, we used either the ff14SB^59^ or the newer ff19SB^30^ protein FFs in conjunction with OL3 or OL3_CP_ to describe the solute. Note that ff19SB was directly parametrized to be used with the OPC water model.^30^ To test the TIP4PD water model, we have used the DES-Amber FF for the RNA^38^ and the protein.^60^

### System building

All simulated systems except the DES-Amber simulations were first solvated in an octahedral box filled with water molecules, ensuring a minimum of 12 Å between any solute atom and the box edge. Unless specified otherwise, an excess KCl salt concentration of 0.15 M was achieved by randomly placing KCl ions around the solute; see below a separate paragraph detailing all the water models and ion parameters we tested. Energy minimization and equilibration procedures were performed with pmemd.MPI in AMBER 22, following the established protocol.^61^ Production simulations were conducted using pmemd.cuda^62^ on RTX 3080ti GPUs, with a typical length ranging between 3 to 10 μs. In all cases, multiple independent simulations were initiated with independent equilibration procedures and different random seeds for initial atomic velocities to generate statistically robust ensembles. SHAKE^63^ constraints and hydrogen mass repartitioning^64^ were applied in all simulations, enabling a 4 fs integration time step. Long-range electrostatics were treated with the particle mesh Ewald method^65^ under periodic boundary conditions, with a non-bonded LJ interaction cut-off of 9 Å. Temperature and pressure were maintained using a Langevin thermostat and Monte Carlo barostat, respectively. Because DES-Amber parameters are not currently implemented in AMBER, simulations using the DES-Amber RNA FF combined with the TIP4PD water model were performed in GROMACS2020.^66^ The simulation protocol in GROMACS2020 slightly differed from the one in AMBER 22 due to differences in the simulation codes. Specifically, GROMACS simulations were performed in a rhombic dodecahedral box and bonds involving hydrogens were constrained using the LINCS algorithm.^67^ The cut-off distance for the direct space summation of the electrostatic interactions was 12 Å and the simulations were performed using the stochastic velocity rescale thermostat.^68^

### Water models and ion parameters

For many of the water models tested in this work – TIP3P,^33^ SPC/E,^34^ OPC3,^35^ TIP4PEW^36^ and OPC^19^ – there are multiple possible choices of compatible ion parameters available and their use is not standardized in the literature. We have finally decided to chiefly utilize the combinations currently present in the latest default AMBER source packages of Leap^58^ for each water model (i.e. leaprc.water.*; where * is the water model), as we expect this is how majority of people will be performing their MD simulations. This gave us the standard combinations of Joung&Cheatham (JC) KCl parameters^69^ to be used with the TIP3P, SPC/E and TIP4PEW water models and Li&Merz (LM) parameters^70-71^ for the OPC and OPC3 water models. Nevertheless, limited cross-testing of these parameters was also carried out for selected systems to verify the potential influence of the ion parameters on the observed structural transitions. The simulations strongly suggest that the choice of monovalent ion parameters does not have any effect on the properties investigated in the present study. For the magnesium ions, the 6-12 Li&Merz parameters^72^ were utilized in all cases. Note that not all the water models were tested for every system. DES-Amber simulations with the TIP4PD water model were always run with the scaled CHARMM22^73^ ion parameters.^38^

### REST2 enhanced sampling simulations of the hTTR system

Compared to the other systems, the hTTR showed slower conformational developments (see below). In addition to the standard MD simulations of the hTTR system, we have therefore also performed REST2^74^ enhance sampling simulations. The number of replicas for the calculations done in AMBER was 7, with the scaling factors ranging from 1 to ∼0.6 and average successful exchange rates of ∼25%. The hot zone was defined in order to support the sampling of the opening of the TTR interaction. Specifically, the base pairs^75^ *t*HW A6:U36 and *c*WW G8:C35 (i.e. the four nucleotides constituting the receptor part of the TTR; original PDB numbering used) except the phosphates were placed in the hot-zone. At the same time, a stabilizing structure-specific HBfix (sHBfix)^76^ of 2 kcal/mol between the hydrogens and acceptors was established to stabilize the H-bonds constituting these base pairs. The sHBfix was not scaled along with the replicas. This setup ensured the REST2 scaling effectively enhances sampling of the tertiary interactions that the receptor forms with the GAAA TL while the sHBfix prevents disruptions of the base pairing H-bonds of the base pairs within the hot-zone. See Supporting Information Text and Figure S2 for further explanation and visualization of the hot zone. All REST2 simulations were carried out with the pmemd.cuda.MPI module in AMBER 22 for 2 μs. The simulation parameters matched those of the standard simulations described above, with the exception that the production phase of the REST2 simulations was conducted in the constant volume (NVT; canonical) ensemble. The REST2 simulations using the DES-Amber FF and TIP4PD water model were performed in GROMACS2018 in combination with PLUMED2.5^77^ using the Hamiltonian replica exchange implementation.^78^ The REST2 GROMACS protocol matched the hot-zone definition and the application of the stabilizing sHBfix potential used in the AMBER calculations. However, due to technical limitations prohibiting odd number of replicas, eight replicas had to be used. In this case, the scaling factor (λ) values ranged from ∼1.07 to ∼0.60, with the second replica serving as the reference (scaling factor 1; unbiased), with an exchange rate of approximately 30% throughout the ladder. An additional eighth replica was placed at the bottom rather than the top of the ladder to avoid biasing TTR stability more in GROMACS simulations than in those performed in AMBER.

### Simulations with mixed water boxes

To directly estimate the affinity of the different water models to the RNA solute atoms, we constructed systems with mixed water boxes containing equimolar amounts of waters of different models. In this fashion, we constructed three different systems, each containing a cytidine nucleoside surrounded by 1858 OPC–SPC/E, 1970 OPC3–SPC/E and 1982 TIP4PD–SPC/E water molecules, respectively. To allow the comparison, the OL3 FF was utilized in each case to describe the nucleoside. The systems were built by first adding the corresponding amounts of OPC, OPC3 and TIP4PD waters in xLeap. This was followed by using parmed to turn exactly half of the waters into the SPC/E parametrizaton, by altering the parameters and deleting superfluous atoms if present. We note this building approach already creates relatively randomized water mixtures which get further rapidly mixed during the subsequent equilibration phase. Three standard MD simulations were performed for each system, and we monitored the populations of H-bonds formed between the two water models and the H-bond donors and acceptors of the cytidine nucleoside.

### Analyses

Analyses and visualization of the MD trajectories were carried out using cpptraj and VMD software packages.^79-80^ Graphs were generated with gnuplot, while molecular renderings were created using povray. Conformational transitions in the simulated systems were primarily identified through visual inspection of the simulation trajectories, since we monitored large conformational changes (disruption of tertiary interactions). In most cases, the structural changes were sufficiently extensive and visually prominent that standard RMSD analysis relatively to the experimental structure was fully adequate to capture the differences observed for the different water models. Additional analysis method was evaluation of the presence of selected tertiary H-bonds based on the interatomic distance and angle of the donor and acceptor groups. Distances between heavy atoms under 4.0 Å and donor-hydrogen-acceptor angles above 120° were considered as cutoff for the H-bonds. Analyses of the REST2 trajectories were performed on the reference (unscaled) replica, as well as on demuxed continuous trajectories. The protein-RNA interface size in the L1 stalk system was calculated by subtracting the solvent-accessible surface area^81^ of the complex from the sum of the individual RNA and protein surface areas.

Analysis of native protein-RNA contacts in the was performed by defining heavy-atom pairs within 6 Å in the starting structure as native.

## Results and discussion

### Basic Overview of the Simulations

Below we present results of nearly half a millisecond of MD simulations, comparing the performance of different water models for three RNA systems stabilized by long-range tertiary interactions: two variants of the L1 stalk ribosomal RNA (rRNA), one of them also with the bound L1 protein, the miniTTR-6 construct with a specific fold and one TTR interaction, and the hTTR duplex homodimer with two symmetrical TTR interactions (see Methods and Table 1). The primary metric used for evaluation was the preservation of native RNA folds observed in the experimental structures.

**Table 1.**
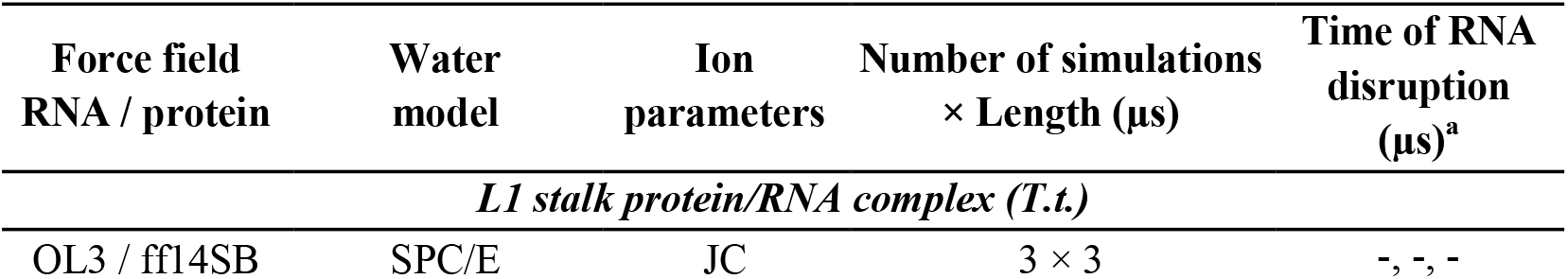
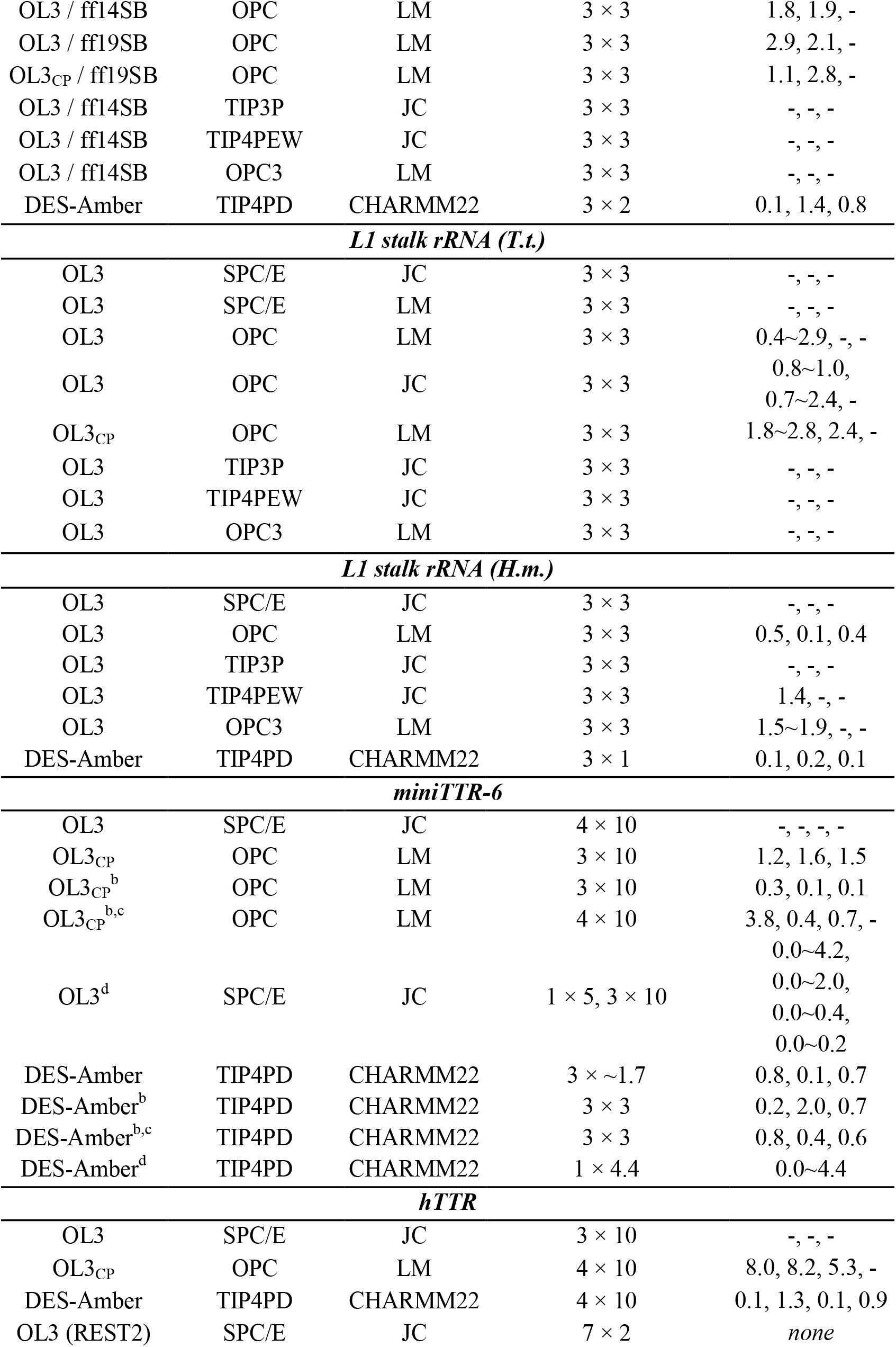
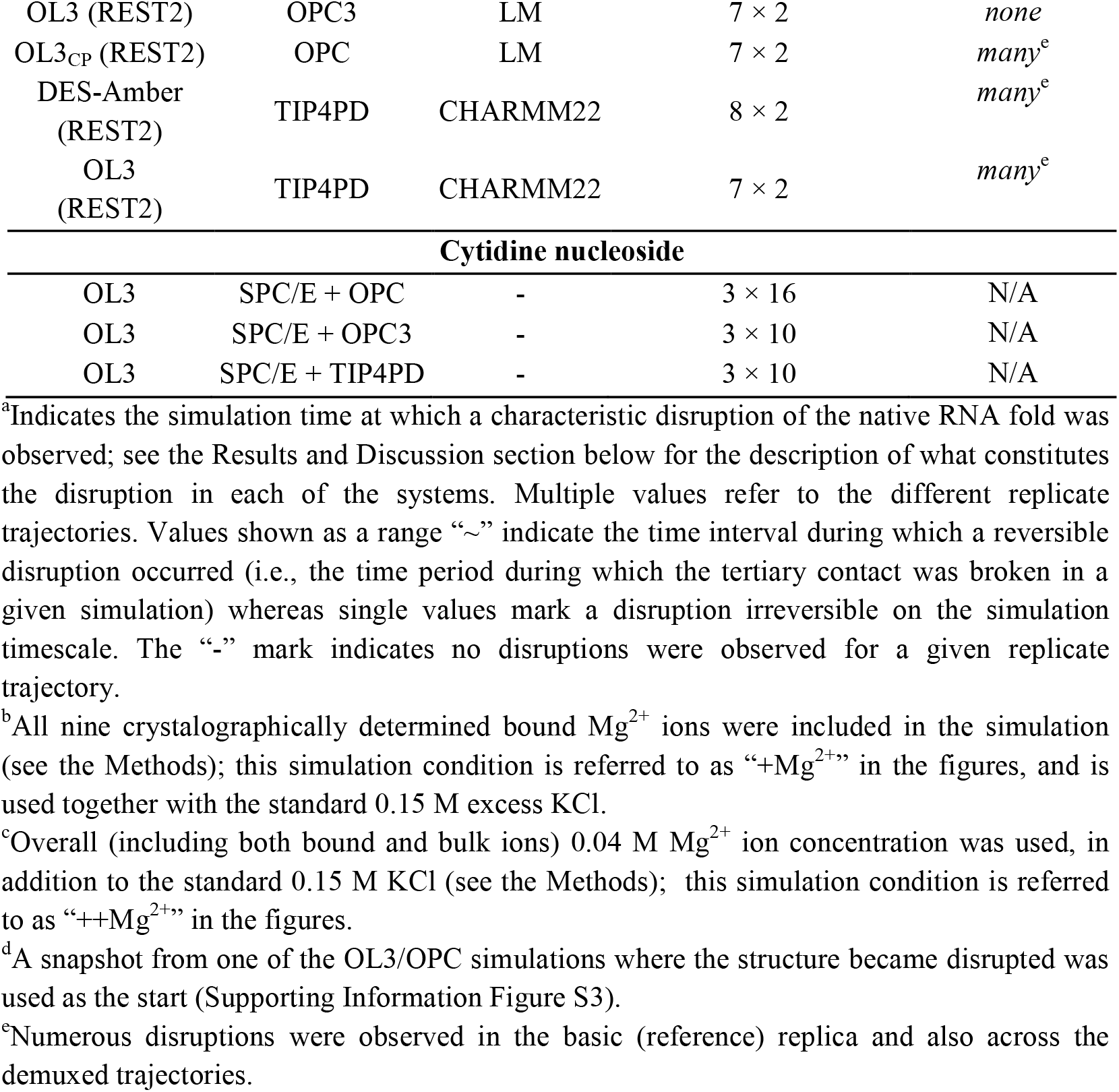
List of MD simulations.

For all systems, the four-point OPC and TIP4PD water models were universally less successful in maintaining the native folds, with both reversible and irreversible disruptions observed that weren’t seen with the other tested water models. We note that use of the adjusted phosphate van der Waals (vdW) parameters by Steinbrecher et al.^20^ along with the OL3 FF did not prevent the loss of the native RNA folds in the OPC simulations. In other words, the OL3 and OL3_CP_ RNA FF variants provided qualitatively identical performance within the limits of our sampling (Table 1) and they will be henceforth not described separately. Likewise, cross-testing of the JC and LM monovalent ion parameter (see Methods) sets showed that they did not affect either the loss of native RNA folds in the OPC simulations or the generally good performance observed in the SPC/E simulations (Table 1). In conclusion, the OPC water model appears to be the sole factor responsible for the structural instabilities in the OL3/OPC simulations.

Qualitatively identical instabilities to those seen in the OL3/OPC simulations were also observed with DES-Amber/TIP4PD FF combination. One notable difference came from the apparent tendency of the DES-Amber FF to favor extended A-form-like structures. Namely, once the TIP4PD water model initially “loosened” the native fold, additional structural rearrangements immediately followed, driving the structure even further away from the native fold and preventing its restoration. In contrast to the OL3/OPC simulations, where disruptions were sometimes reversible and the native fold could at least temporarily recover, no such restorations were ever observed with DES-Amber/TIP4PD on our simulation timescale. We suggest that this outcome is primarily attributable to genuine differences between DES-Amber and OL3 RNA FFs; in a recent study we noted that DES-Amber FF (as well as its predecessor DESRES^82^) are somewhat biased in favor of the A-form-like RNA structures.^18^ However, when focusing solely on the structural influence of the water models (the topic of this paper), TIP4PD and OPC exert essentially identical destabilizing effects on the native RNA folds.

### L1 rRNA Undergoes Structural Disruption in OPC Simulations

Isolated L1 rRNA from both *T.t*. and *H.m*. as well as the L1 stalk protein-RNA complex from *T.t*. exhibited reduced structural stability when simulated with the OPC water model (Figure 1 and Table 1). Among the three structures, the isolated L1 stalk rRNA from *H.m*. underwent the most rapid structural degradation, with permanent loss of its structure in entirely all OPC simulations as well as even in one simulation with the TIP4PEW model (an older four-point water model from 2004). The L1 stalk rRNA from *T.t*., whether in isolation or complexed with the L1 protein, was more resistant to structural changes, showing disruptions in only some OPC simulations. In the simulations of the isolated L1 stalk rRNA, the disruptions were additionally reversible on the simulation timescale with one exception (Table 1). In contrast, the DES-Amber/TIP4PD simulations exhibited rapid and irreversible disruption of the L1 stalk in both the isolated rRNA and the protein-RNA complex. Moreover, the structural rearrangements were substantially more extensive (Figure 1C, D), in some cases necessitating early termination of the simulations to avoid solute image clashes. It should be noted that in an earlier study,^38^ stabilization of the L1 stalk structure in DES-Amber/TIP4PD simulation was achieved through the inclusion of very high concentration of bound (experimental X-ray) and bulk Mg^2+^ ions, generally mimicking the conditions used for crystallization.

**Figure 1.**
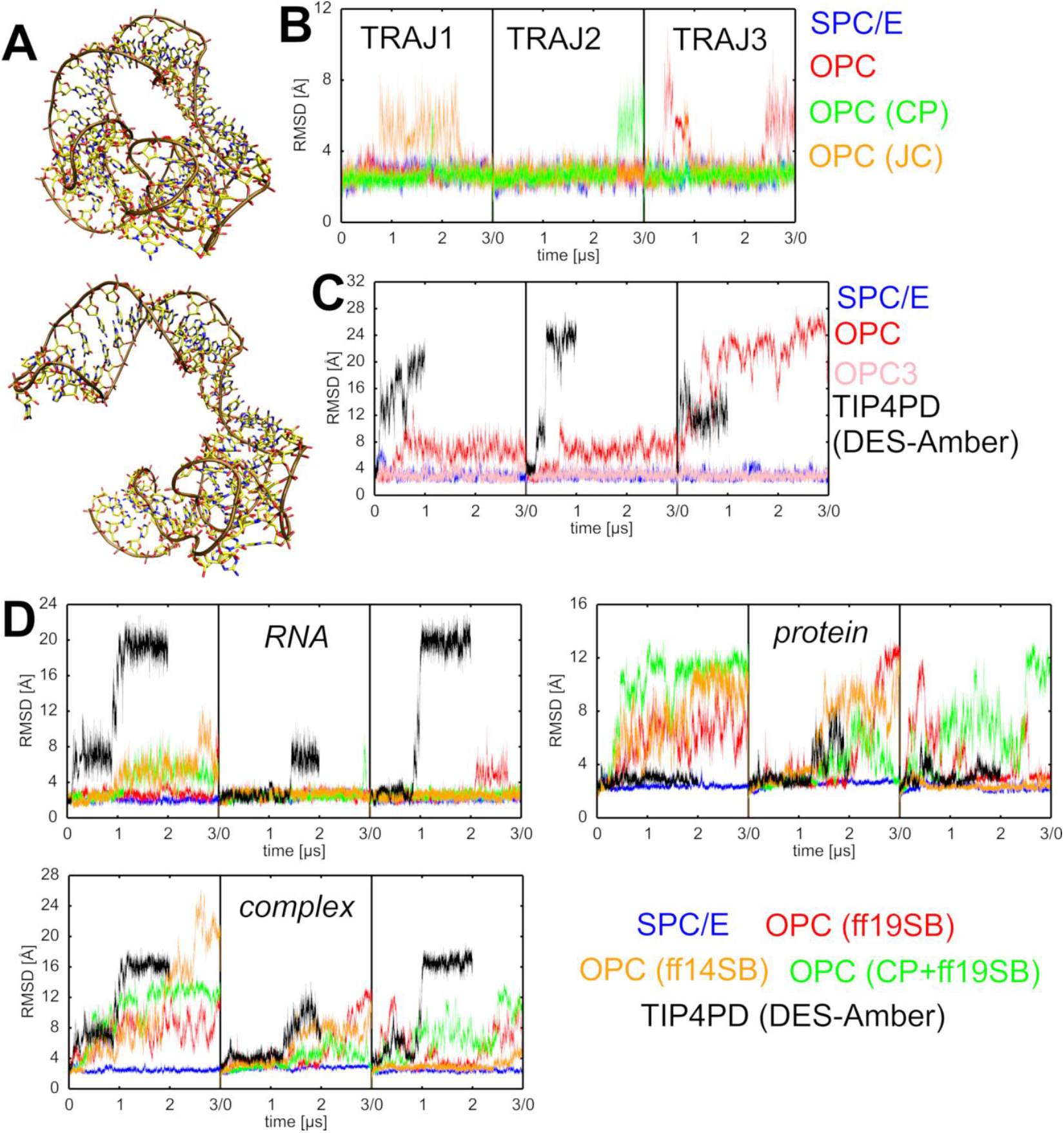
Loss of the native RNA fold in simulations of L1 stalk. **A)** Comparison of the experimental structure (top) and an example of a disrupted structure typically seen in OPC and TIP4PD simulations (bottom). More extended A-form-like structures could be observed with the DES-Amber/TIP4PD FF combination (Supporting Information Figure S4). **B)** Time evolution of the RMSD of the isolated L1 stalk rRNA from *T.t*. in individual simulations using selected water models. Datasets are color-coded according to the legend on the right (see also Table 1). For space saving reasons, all three independent parallel simulations (replicates) per water model are shown in a single plot, with individual trajectories separated by black vertical bars. The time evolution graphs will be presented in this manner henceforth. **C)** Same as B, but for the isolated L1 stalk rRNA from *H.m*. **D)** Time evolution of the RMSD of the L1 stalk protein-RNA complex from *T.t*. Separate RMSD plots are shown for the RNA, the protein, and the entire complex.

Overall, the three-point water models showed superior stability for the L1 stalk system, with merely one short reversible disruption observed for the *H.m*. system (Table 1).

### Collapse of the Protein–RNA Interface of the L1 Stalk in OPC Water Simulations

In addition to the rRNA disruptions described above, we consistently observed a gradual major loss of the protein-RNA interface in the OPC water model simulations (Figure 2). In some of the trajectories, the L1 protein almost entirely disengaged from the rRNA by the simulation end. Notably, these changes appeared to be independent from the RNA disruptions (see above) and were even faster, suggesting a distinct and separate issue attributable to the water model (Figure 2B). The primary loss of the RNA structure occurs rather far from protein binding area. To assess whether these interface effects could be related to the utilized protein FF, we tested both the ff14SB and ff19SB protein FFs. We observed the loss of the interface occurring with both FFs in the same manner. Note that the ff19SB is primarily recommended to be used with the OPC water model by the authors.^30^ The protein-RNA interface is unstable also with the DES-Amber/TIP4PD FF combination (Figure 2B). In contrast, simulations using three-point water models such as SPC/E and OPC3 did not reveal any significant destabilization of the protein-RNA interface in the L1 stalk system. percentage of native heavy atom contacts maintained

**Figure 2.**
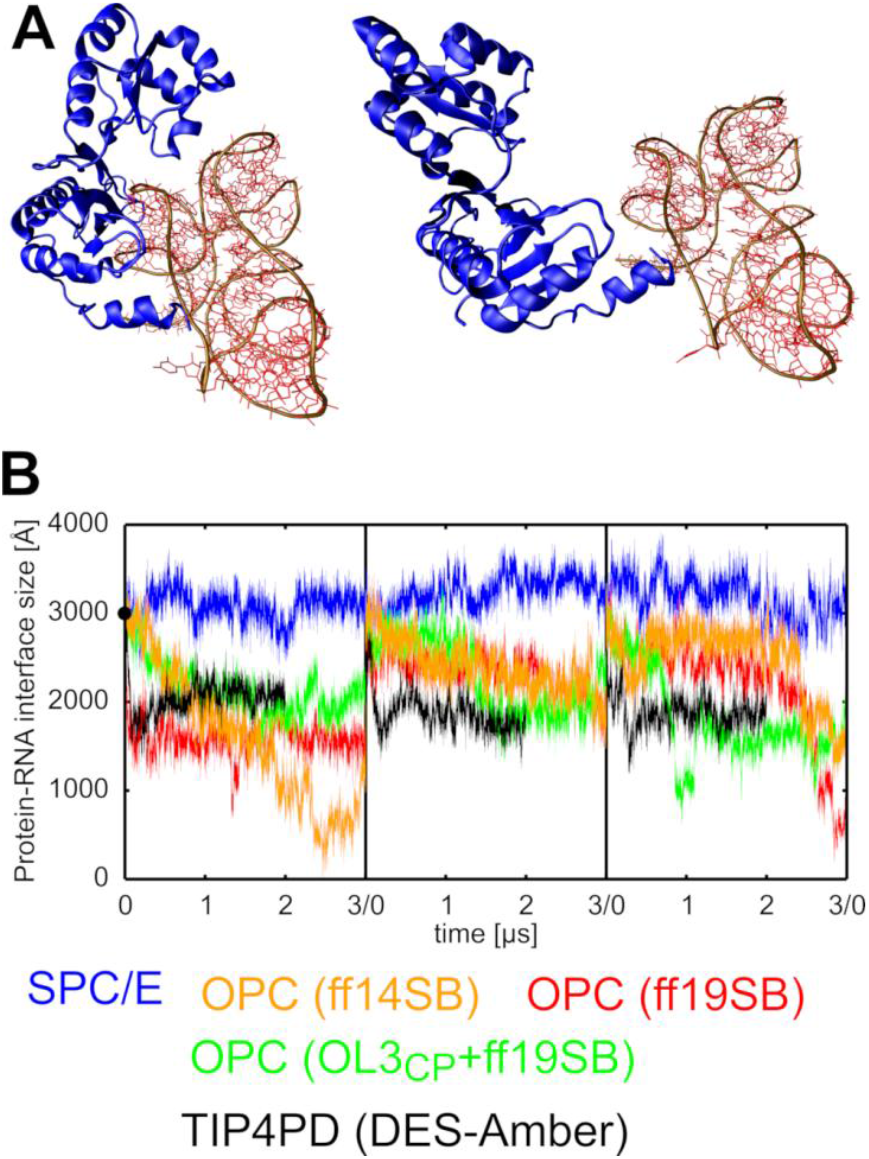
Destabilization of the protein-RNA interface in L1 stalk simulations. **A)** Comparison of the experimental structure (left) and a representative structure from OPC simulations (right), showing the L1 protein nearly fully disengaged from the rRNA. Protein domain is shown as blue ribbons, RNA atoms in red, and the RNA backbone in brown. **B)** Time evolution of the protein-RNA interface size in individual simulations using selected water models. The black circle on the Y-axis indicates the experimental value. Datasets are color-coded according to the legend below the graphs.

### OPC Water Model Induces Large-Scale Unfolding of the miniTTR-6 System

A certain limitation of the L1 stalk system (see above) is that the OPC-induced structural collapse cannot be easily attributed to a well-defined set of problematic solute-solute interactions. Instead, the system appears to undergo diverse and multistep structural rearrangements prior to the structural disruption that furthermore does not occur in every simulation. Nevertheless, among the affected interactions, we noticed those involving RNA 2′-hydroxyl groups as particularly sensitive and usually the first to locally disappear prior to the more global changes. To examine this in a more focused manner, we next selected the miniTTR-6 system, which among other non-canonical structural features (i.e. a kink-turn and IRES element) contains a long-range TTR – a motif known to be stabilized by an extensive network of 2′-hydroxyl-mediated interactions (Figure 3).

**Figure 3.**
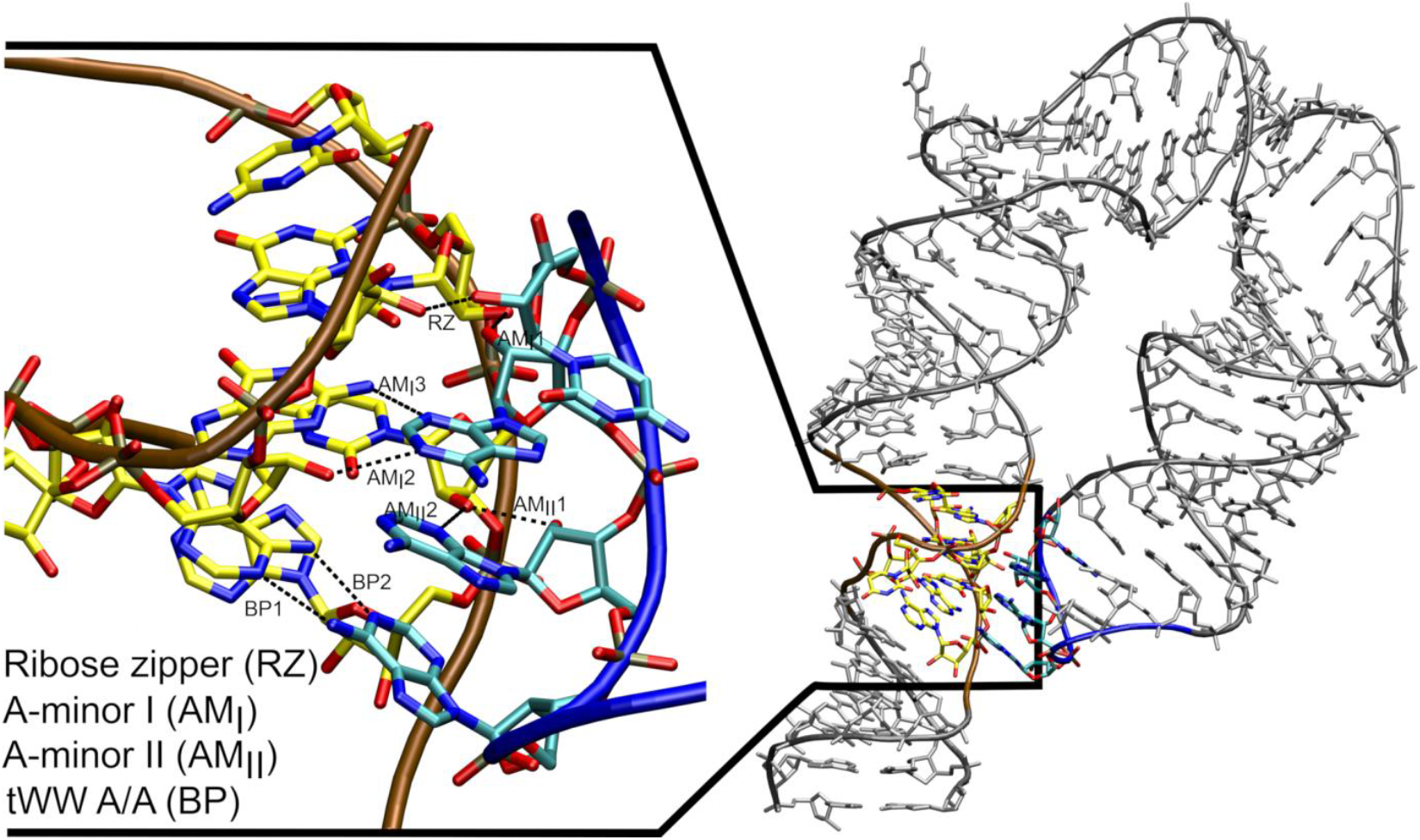
Structural overview of the GAAA tetraloop-tetraloop receptor (TTR) motif within the miniTTR-6. The TTR motif can be divided into four non-canonical interaction submotifs, which are depicted in the Figure. The tetraloop receptor and GAAA TL parts of the motif have their carbon atoms colored in yellow and cyan and their backbone trace in brown and blue, respectively. Individual H-bonds are indicated by black dashed lines and labeled using motif-specific abbreviations. Multiple H-bond interactions within the same submotif are given a numerical suffix to differentiate them. This labeling scheme is applied consistently throughout the present study. Identical TTR motifs are found in the miniTTR-6 and hTTR systems.

As anticipated, simulations of miniTTR-6 using the OPC water model showed rapid disruption of the TTR motif (Figure 4 and Table 1). This was sometimes followed by a straightening of the entire structure. No such disruptions were observed when using the SPC/E water model. We also tested the possible stabilizing influence of the divalent ions by performing the OPC simulations under three different ionic conditions: (1) standard 0.15 M excess KCl concentration, (2) inclusion of the nine crystallographically resolved Mg^2+^ ions (see Methods), and (3) inclusion of both experimental Mg^2+^ ions together with large number of bulk Mg^2+^, to achieve an overall 0.04 M magnesium concentration (see the Methods and Table 1). Both magnesium conditions were supplemented by 0.15 M KCl. However, in all cases, the same TTR disruption was observed (Figure 4 and Table 1), reinforcing the conclusion that the destabilizing effect is primarily attributable to the OPC water model. Notably, when a structure previously disrupted by OPC was re-simulated in a larger SPC/E water box with a standard KCl concentration of 0.15 M, the native fold was fully restored within ∼4.2 μs (Figure 5). We next performed three additional simulations, which also resulted in full restoration in two cases and a partial restoration in the third (Supporting Information Figure S5). These amazing recoveries highlight that the OL3 FF, when paired with SPC/E water, is eventually capable of reversing OPC-induced damage, at least in some cases. Importantly, the result pinpoints the water model as the prime factor in the observed destabilization and indicates that the basic OL3 FF, albeit being far from flawless, provides rather reasonable description of many folded RNAs.^18^

**Figure 4.**
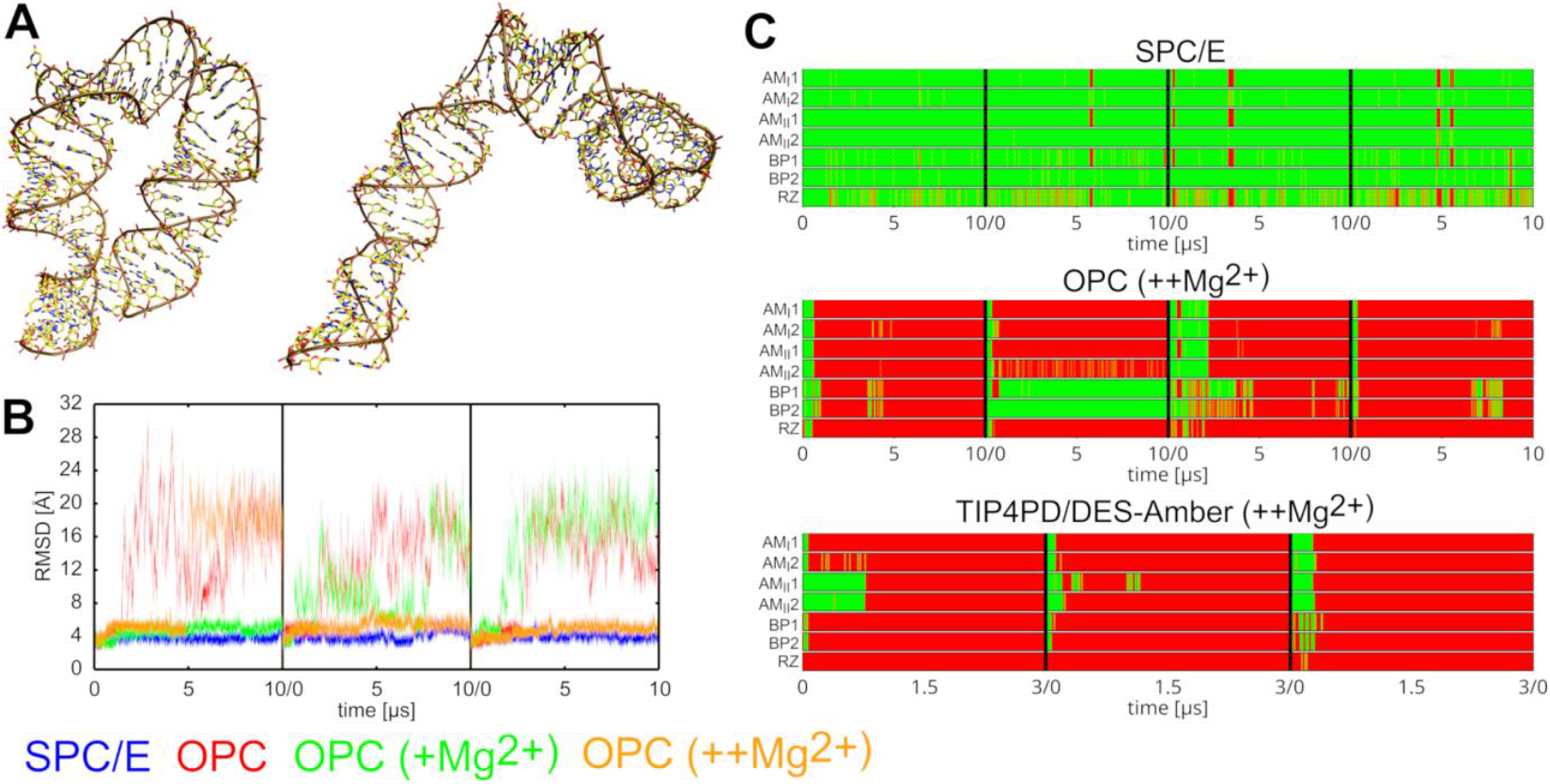
Loss of the native RNA fold in simulations of miniTTR-6. **A)** Comparison of the experimental structure (left) and a selected representative structure from OPC simulations (right). **B)** Time evolution of the RNA RMSD in selected simulations using different water models and ionic conditions (see Methods). Datasets are color-coded according to the legend at the bottom (see the footnotes of Table 1 for explanation of the abbreviations used to define the Mg^2+^ conditions). **C)** Time evolution of the key TTR H-bonds in selected simulations using different water models. Green and red indicate presence and absence of the H-bond, respectively. See Figure 3 for definition of the individual H-bonds. It is notable that the inclusion of Mg^2+^ ions does not stabilize the structure in the simulations. According to the experiments the miniTTR6 structure folding is greatly aided already by very low concentrations of Mg^2+^, but in principle it folds also in excess of monovalents.^52^

**Figure 5.**
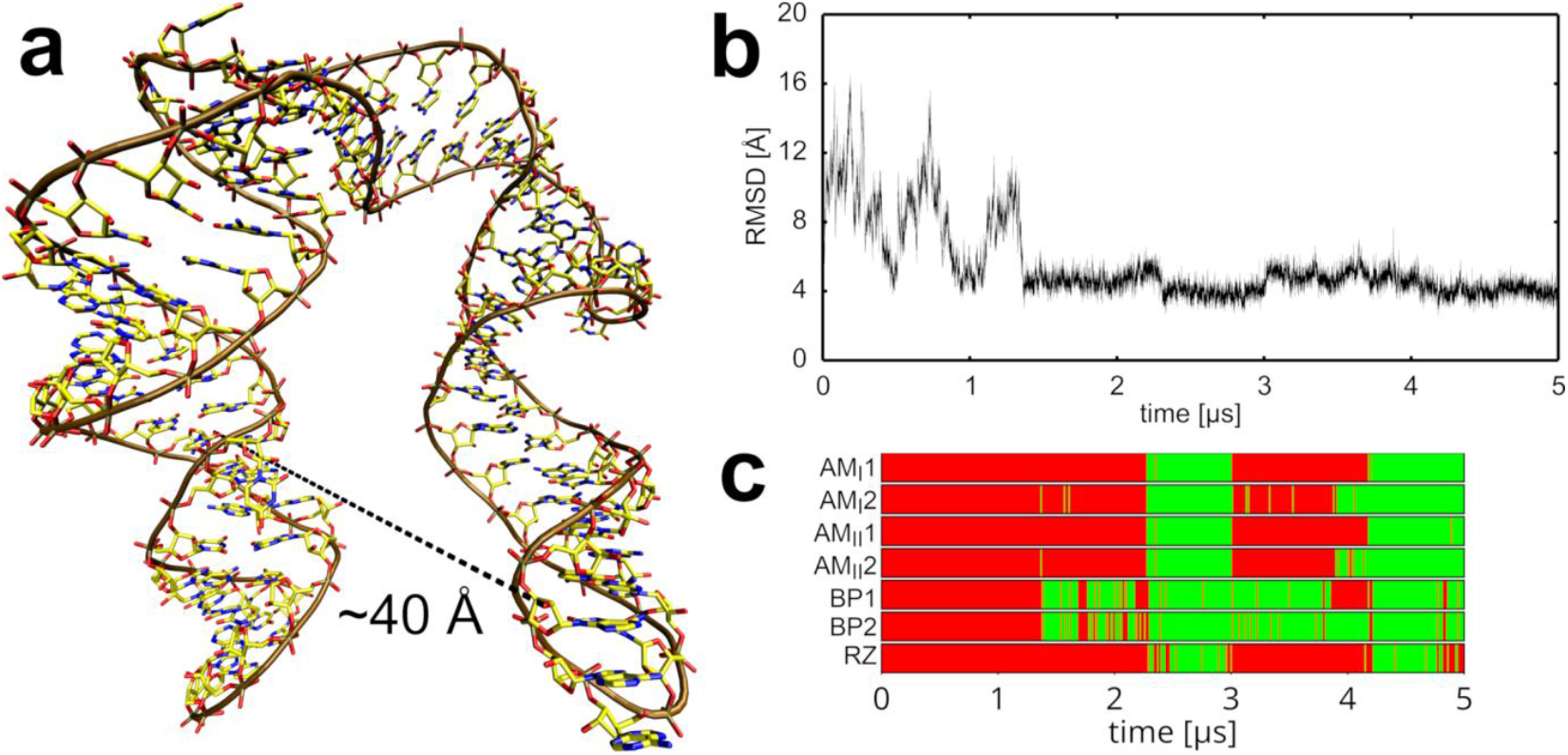
Restoration of the native fold of miniTTR-6 in SPC/E simulation. **A)** A structure previously disrupted in OPC simulations, with the TTR binding partners positioned far apart. **B)** Time evolution of the RNA RMSD in the restoration simulation using the SPC/E water model. **C)** Time evolution of the signature TTR H-bonds. Green and red indicate presence and absence of the H-bond, respectively. See Figure 3 for definition of the individual H-bonds. See also Supporting Information Figure S5 for additional replicates of this.

### DES-Amber/TIP4PD Setup Exhibits Similar Instabilities in miniTTR-6 Simulations as the OL3/OPC combination with an Additional Straightening of the Structure

In our present simulations of the miniTTR-6 system, all the DES-Amber/TIP4PD simulations consistently led to disruption of the TTR motif (Figure 4C and Table 1), followed by extensive global rearrangement (Supporting Information Figure S6). Similarly to the OL3/OPC combination, this occurred regardless of the inclusion of the Mg^2+^. In contrast to OL3, where we showed the OPC-disrupted structures could be restored by transferring them into SPC/E water, the same approach could not be tested with DES-Amber. This is because DES-Amber incorporates solute-solvent NBfix modifications specifically tailored for TIP4PD,^38^ rendering it incompatible with other water models. Nevertheless, we have enlarged the water box to eliminate any image clashes and potentially allow the system to spontaneously relax back to the native conformation. However, no such restoration was observed. Instead, the system exhibited a clear tendency to deviate further from the native fold which indicates potential bias in the RNA FF towards the A-form (Supporting Information Figure S7).

### Simulations of the GAAA Tetraloop-Tetraloop Receptor (hTTR) Homodimer Complex Consistently Reveal a Loss of one TTR

As the final test system, we selected a construct composed of two symmetrical A-RNA helices forming a homodimer via two identical TTR motifs. Unlike the miniTTR-6 system, which contains additional non-canonical motifs that could contribute to the structural changes in some fashion, the hTTR construct effectively isolates the canonical TTR motifs (Figure 3) in a minimal but experimentally well-defined context.^51^ Indeed, in our simulations, the hTTR construct exhibited overall greater structural stability than miniTTR-6, supporting our hypothesis that additional structural strain in miniTTR-6 might be influencing the disruption events. Nevertheless, simulations using the OPC water model still displayed signs of instability. Namely, in three out of four OL3/OPC simulations, one of the two TTRs was irreversibly disrupted (Table 1 and Figure 6) while the second TTR remained stable over the course of the simulation. Likewise, the DES-Amber/TIP4PD simulations revealed loss of one TTR in all four trajectories, on even a shorter timescale than the OL3/OPC (Figure 1). In contrast, no TTR disruptions were observed in simulations using the SPC/E or OPC3 water models, conclusively showing the four-point OPC and TIP4PD water models as the destabilizing factor.

**Figure 6.**
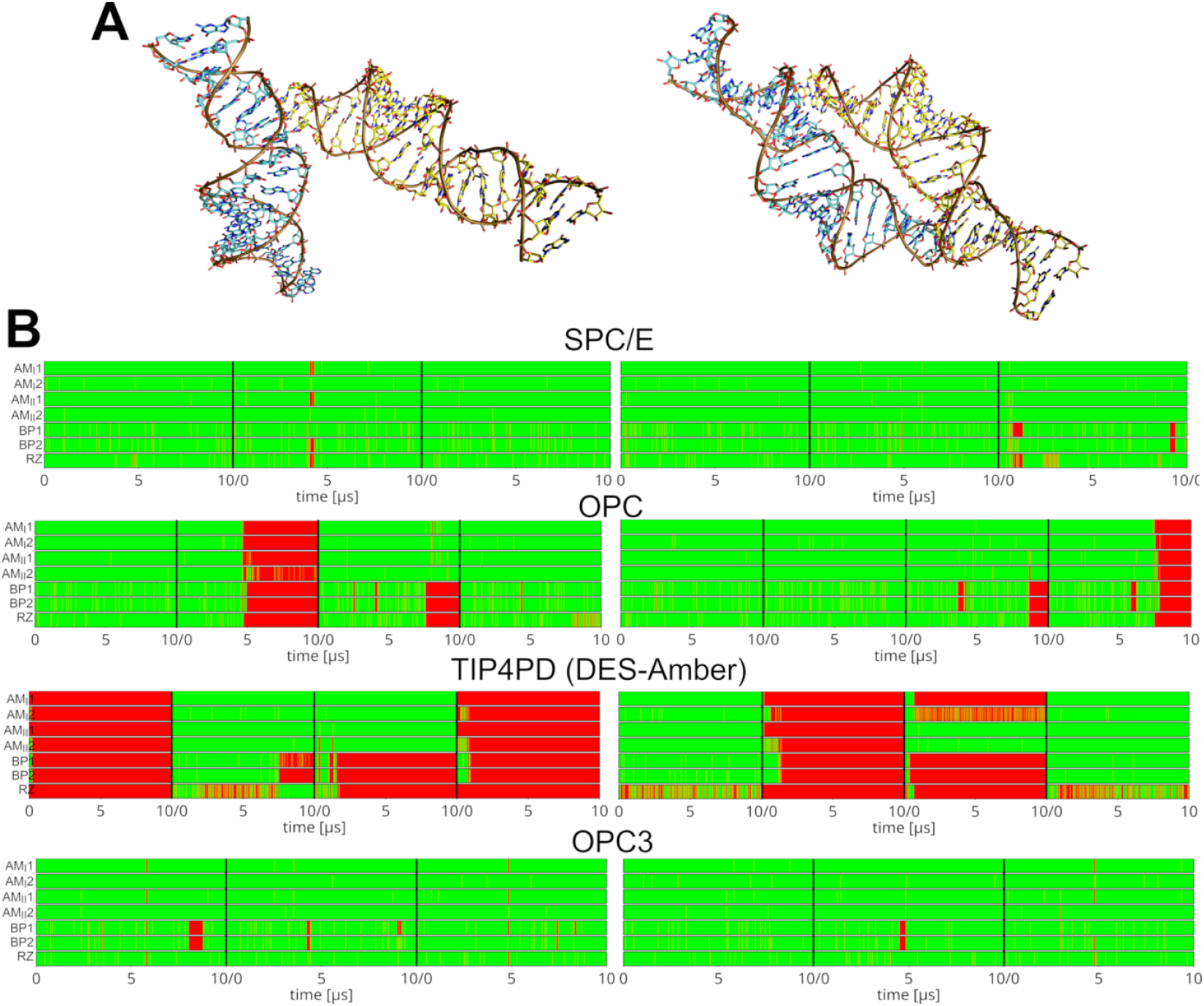
Loss of the native RNA fold in standard MD simulations of the hTTR. **A)** Comparison of the experimental structure (left) and a typical structure observed in simulations using the OPC or TIP4PD water models where only one of the two TTRs remains intact (right). The two helices forming the hTTR have their carbons colored in yellow and cyan, respectively. **B)** Time evolution of the signature TTR H-bonds (see Figure 3) in simulations using different water models. Green and red indicate presence and absence of the H-bond, respectively. See Figure 3 for definition of the individual H-bonds. The left and right graphs show the first and second TTR site in the hTTR system, respectively.

### REST2 Enhanced Sampling Simulations Confirm OPC- and TIP4PD-Induced Destabilization of TTR

In standard MD simulations of the hTTR system using the OPC or TIP4PD water models, only one out of the two TTR motifs was disrupted on the timescale of our simulations (see above and Figure 6). In addition, for the OL3/OPC combination, the disruption events occurred relatively late in the simulations. Nevertheless, the inferior performance of OPC and TIP4PD was evident, even though a fully converged picture was not obtained.

To further examine this, we next performed REST2 enhanced sampling simulations of the hTTR using the SPC/E, OPC, OPC3, and TIP4PD water models (Figure 7; see also Methods and Table 2). We observed that in the reference (unbiased) replica, the SPC/E and OPC3 models maintained almost fully stable TTR motifs throughout. Even in higher replicas there were only brief fluctuations. In contrast, the OPC REST2 simulations showed quick onset of TTR instability (Figure 7), including a rare simultaneous loss of both TTR motifs in some continuous trajectories (Supporting Information Figure S8). Note that in standard MD simulations, we could always observe only loss of a single TTR motif. All the disruptions observed in the OPC REST2 simulations were fully reversible on the simulation timescale as the system was intermittently switching between the disrupted and bound states at the two ends of the homodimer. Even when both TTR motifs were transiently lost and the two helices briefly drifted apart randomly in the box, the native interface was eventually reformed over time (Supporting Information Figure S8). We suggest this conclusively shows the OPC-induced destabilization of the hTTR structure while also demonstrating the good performance of the OL3 RNA FF for the TTR interaction. This result contrasts sharply with the DES-Amber/TIP4PD REST2 simulations, which showed only irreversible losses of the TTR, with simultaneous loss of both TTRs having occurred in over half of the continuous trajectories by the simulation end. To disentangle the effect of the solute force field from that of the water model, we performed additional REST2 simulations with the non-standard OL3/TIP4PD combination (Table 1). In this setup, performance for the hTTR improved markedly, still remaining somewhat inferior to OL3/OPC, but comparable (Figure 7 and Table 2).

**Table 2.** Populations (in %) of conformational states sampled in REST2 simulations of the hTTR.^a^.

| Water model / Conformation | Both TTRs stable | Single TTR stable | No TTR stable |
| --- | --- | --- | --- |
| **SPC/E** | 100 | 0 | 0 |
| **OPC3** | 96 | 4 | 0 |
| **OPC** | 62 | 38 | 0 |
| **DES-Amber/TIP4PD** | 0 | 75 | 25 |
| **OL3/TIP4PD** | 51 | 34 | 15 |

<sup>a</sup>The hTTR system contains two TTR motifs, either of which could become disrupted in the REST2 simulations. Reported values correspond to the last 500 ns of the unscaled (basic) replica. The presence of a TTR interaction was evaluated by measuring the distance between the centers of geometry defined by the tetraloop and tetraloop receptor H-bonding heavy atoms, respectively. Distances between the centers greater than 12 Å were classified as disrupted motifs (i.e. the TTR not present). By measuring this distance for both TTR motifs, we classified the ensemble states into three categories: both TTRs present, only one TTR present (either of them), or no TTR present.

**Figure 7.**
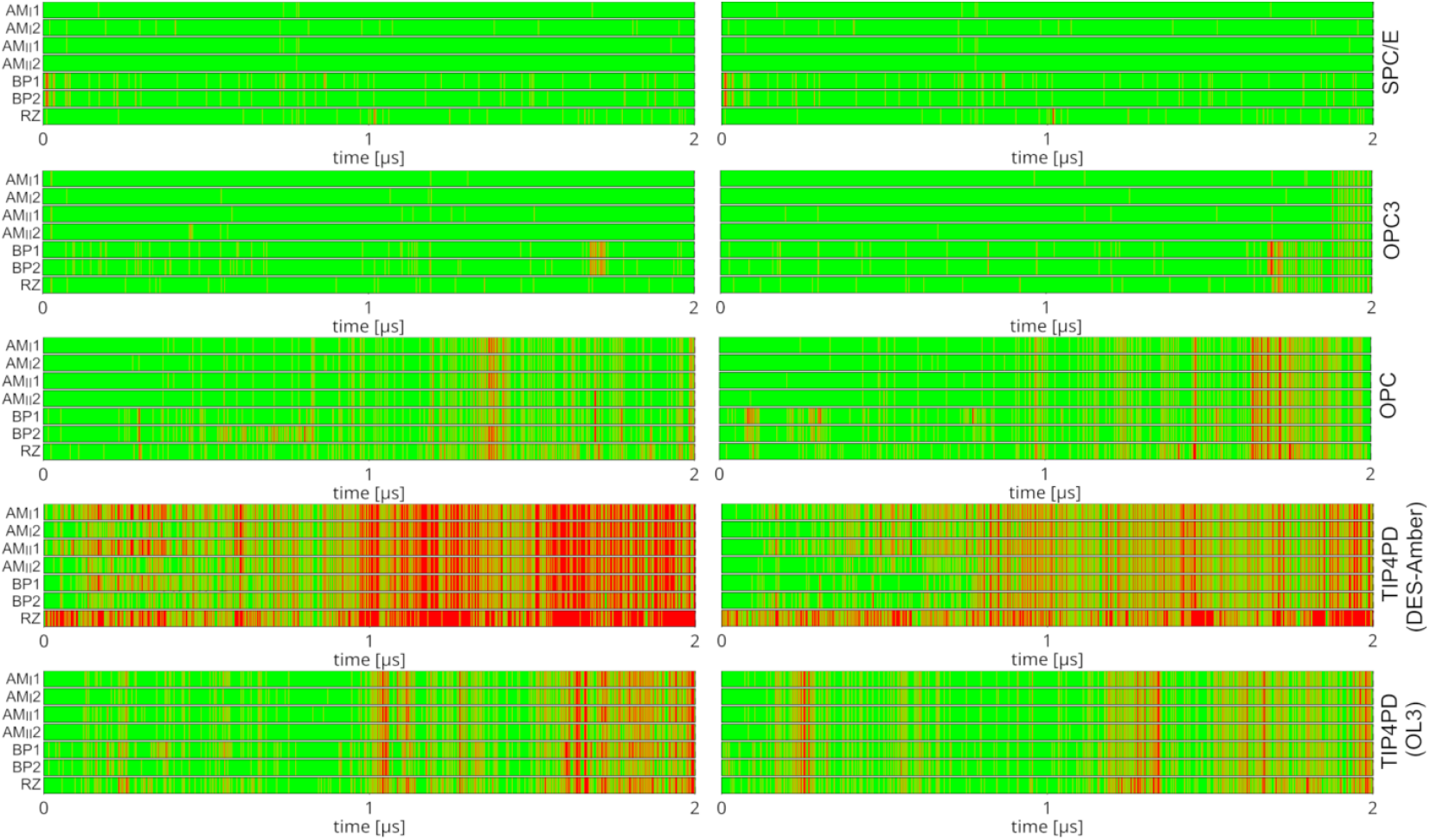
REST2 enhanced sampling simulations of the hTTR. A) Time evolution of the signature TTR H-bonds in the unscaled replica of REST2 simulations using different water models. Green and red indicate the presence and absence of each hydrogen bond, respectively. See Figure 3 for definitions of individual interactions. Left and right graphs show the first and second TTR site in the hTTR system, respectively.

### OPC and TIP4PD Waters Preferentially Interact with RNA H-bond Donor/Acceptor Groups Relative to SPC/E

Detailed inspection of all trajectories of the tested systems indicate that the OPC and TIP4PD waters destabilize the RNA structures by excessively competing against native RNA–RNA hydrogen bonding. In other words, H-bond donors and acceptors in RNA seem to form more frequent interactions with the tested four-point water models compared to the three-point models (data not shown), which might explain the observed disruptions of the native intramolecular interactions essential for the RNA structural integrity. While the trend is qualitatively clear, quantifying the extent of this “water insertion” interference for the large systems is challenging due to sampling limitations, presence of numerous such interactions and complexity of the conformational changes. In attempt to provide more quantitative insights, we constructed model systems consisting of just a single cytidine nucleoside solvated in an equimolar mixture of two different water models (see Methods). We then calculated the relative populations with which the different water models interact with the individual H-bond donors and acceptors of the nucleoside in standard simulation. The rationale is that the relative occupancies of each water model at specific RNA donor or acceptor sites should reflect the relative affinities of the water model to these RNA sites. Any deviation from the 1:1 binding ratio indicates preferential binding by one water model to the site. The free energy difference can then be estimated using the Boltzmann two-state model from the relative populations (Table 3) and related to stability of the respective RNA-RNA H-bonds in a given water model.

**Table 3.** Difference in estimated binding free energy (in kcal/mol) between the specified water molecules and the specific H-bond donors and acceptors of the cytidine nucleoside.^a^.

| RNA atom | OPC – SPC/E | OPC3 – SPC/E | TIP4PD – SPC/E |
| --- | --- | --- | --- |
| **acceptors** |  |  |  |
| **O5'** | 0.05 | -0.03 | -0.01 |
| **O2'** | 0.08 | -0.05 | -0.04 |
| **O3'** | 0.07 | -0.04 | 0.00 |
| **O2** | 0.12 | -0.01 | 0.10 |
| **N3** | 0.13 | -0.02 | 0.04 |
| **donors** |  |  |  |
| **HO5'** | 0.11 | 0.04 | 0.18 |
| **HO2'** | 0.10 | 0.01 | 0.16 |
| **HO3'** | 0.10 | 0.03 | 0.14 |
| **NH41** | 0.02 | -0.01 | 0.15 |
| **NH42** | 0.01 | 0.01 | 0.18 |

<sup>a</sup>Negative number means the RNA donor/acceptor interacts more strongly with the SPC/E water. Values represent a combined simulation ensemble of all three simulations. Virtually no variance was observed among simulation replicates after $\sim 2 \mu\text{s}$ , indicating excellent convergence. A nucleoside was used for the simulations to avoid the need to include a counter-ion.

Across three independent trajectories, we observed the OPC water molecules interacting more strongly with both RNA donor and acceptor atoms compared to the SPC/E water model. It was particularly visible with the 2′-hydroxyl groups and base atoms N3 and O2 (Table 3). A slightly different picture was observed for the TIP4PD, namely, for some donor groups this water model interacted less efficiently than the SPC/E. However, this was compensated by much stronger interactions with the acceptors, leading to similar expected net shift of the RNA-RNA H-bond stability as for the OPC water (Table 3). Finally, we note that the OPC3 water model has reduced and increased affinity to donors and acceptors, respectively, compared to the SPC/E model. This might be resulting in only marginal stability changes for the RNA-RNA H-bonds, in agreement with the RNA simulations. While the differences per single H-bond interaction shown in Table 3 are quite small, they may become significant for large RNAs where many such interactions form and collectively contribute to the stability of the native RNA folds.

## Conclusions

Through an extensive series of MD simulations on a diverse set of structured RNA systems, we have demonstrated that the OPC and TIP4PD water models can induce structural instability in folded RNAs. This behavior was observed across multiple RNA systems of varying size and complexity, including the ribosomal L1 stalk, miniTTR-6 construct, and GAAA tetraloop-tetraloop receptor homodimer (hTTR) complex, using a selection of popularly used RNA FFs and ion parameters. In each case, the water model was identified as the primary factor responsible for the observed structural instability of the RNA fold, independent of the RNA FF, ion parameters, or ion concentration. These findings raise some concerns about the increasingly common use of the OPC and TIP4PD water models in RNA simulations, particularly in systems involving complex hydrogen-bonding networks and tertiary interactions. The problem extends also to protein-RNA interfaces as complete loss of the L1 stalk complex interface was observed in some simulations using the OPC water model. Analysis of water occupancy around RNA donor and acceptor atoms revealed that OPC and TIP4PD water molecules interact with the RNA with greater affinity than SPC/E water, by ∼0.1-0.2 kcal/mol per hydrogen bond. We suggest that this introduces direct competition with the native hydrogen-bonding patterns and ultimately destabilizes RNA folds. In small systems such as RNA tetranucleotides or hairpins, this effect can be negligible or even beneficial, as it tends to favor less compact conformations. However, in larger folded RNAs, it may introduce a significant energetic penalty that undermines the formation and stability of the native fold. Particularly affected are motifs where the donors/acceptors are solvent-exposed or structurally flexible, such as the 2′ hydroxyl groups.

Taking together, our results indicate that the OPC and TIP4PD water models, despite their favorable bulk properties and recent popularity in MD simulations, may not be best suited for maintaining the structural integrity of some folded RNAs or even protein-RNA complexes. Their tendency to over-solvate hydrogen bonding groups undermines native interactions and leads to destabilization that cannot be mitigated by choice of RNA FFs or ion model selection. For folded RNA structures and protein-RNA complexes, the three-point water models such as SPC/E, TIP3P and OPC3 might be more reasonable choices.

## Funding

This work was supported by the Czech Science Foundation (grant number 23-05639S; to M.K. V.M., and J.Š.). P.B. and M.O. were supported by ERDF/ESF project TECHSCALE (No. CZ.02.01.01/00/22_008/0004587). M.O. also acknowledges the financial support of the European Union under the REFRESH – Research Excellence For REgion Sustainability and High-tech Industries project number CZ.10.03.01/00/22_003/0000048 via the Operational program Just Transition. This research also received the support of EXA4MIND, a European Union’s Horizon Europe Research and Innovation programme under grant agreement N° 101092944 (V.M., P.B. and M.O.). Views and opinions expressed are however those of the author(s) only and do not necessarily reflect those of the European Union or the European Commission. Neither the European Union nor the granting authority can be held responsible for them.

## Acknowledgement

This work has been conducted in the sustainability period of the project SYMBIT No. CZ.02.1.01/0.0/0.0/15_003/0000477 as its follow-up activity. We acknowledge the use of CESNET data storage facilities [grant number LM2018140].

## Supporting Information

Additional details of the REST2 protocol utilized in simulations of the hTTR. Supporting Figures and Tables.

## Notes

### Competing Interest Statement

The authors have declared no competing interest.

### Summary of Updates

We have included additional data relevant to the study. As a result, two additional authors were also added to the author list.

