## Supporting Information for "Destabilization of Structured RNAs by OPC and TIP4PD Water Models"

### Supporting Information Text

**Additional comments on the definition of the hot zone region in the REST2 simulations of the hTTR system.** The two base pairs of the tetraloop receptor (A6:U36 and G8:C35) we have chosen to be included in the list of hot zone (scaled) atoms form two distinct types of H-bonds: (**a**) the standard base pairing H-bonds and (**b**) tertiary H-bonds with the GAAA tetraloop which constitute the TTR motif (Figure S2). Note that in this REST2 setup, (**a**) are directly scaled by $\lambda$, while (**b**) are scaled as $\sqrt{\lambda}$ as the GAAA tetraloop lies outside the hot zone. Since our goal was actually to only accelerate the sampling of (**b**), we applied additional stabilization using 2 kcal/mol sHbfix potentials for every base pairing H-bond in (**a**) to compensate for their weakening. This effectively focused the sampling enhancement onto the tetraloop-tetraloop receptor (TTR) interactions (**b**) (Figure S2). Note that we deliberately avoided including the GAAA tetraloop in the hot zone as this would have shifted most of the sampling enhancement toward exploring conformational variations within the GAAA tetraloop rather than the tertiary interactions forming the TTR motif. Such variations were not the target of this study. In addition, these interactions are less straightforward to stabilize with sHbfix restraints, making the two tetraloop receptor base pairs a more effective choice to define the hot zone.

**Supplementary Information Figures**

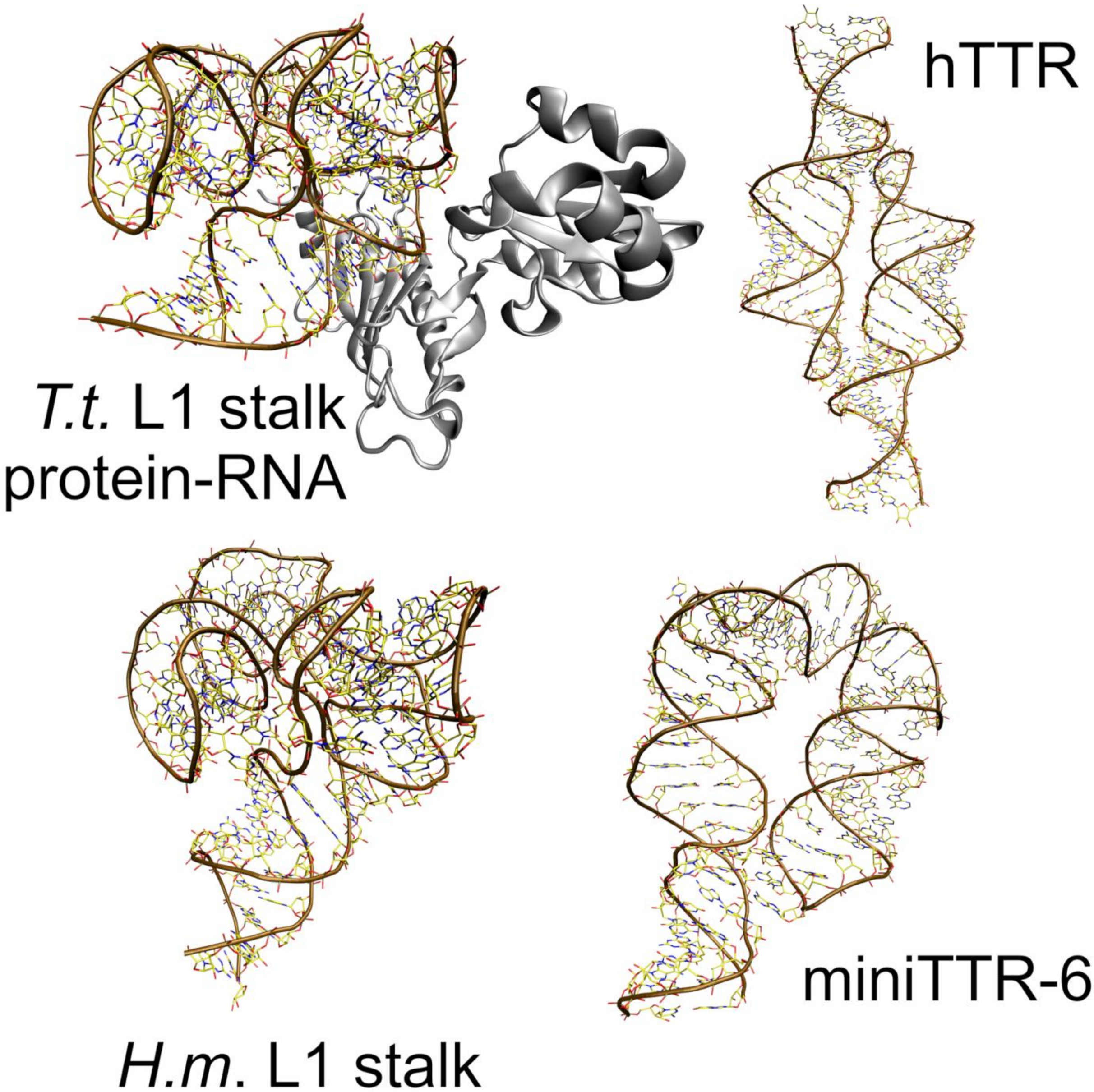


Figure S1. **Structured RNAs and protein-RNA complex analyzed in this study.** The T. thermophilus L1 stalk rRNA was also simulated in isolation. RNA atoms are depicted as sticks with the backbone highlighted as a brown tube, while the protein is shown as grey ribbons.

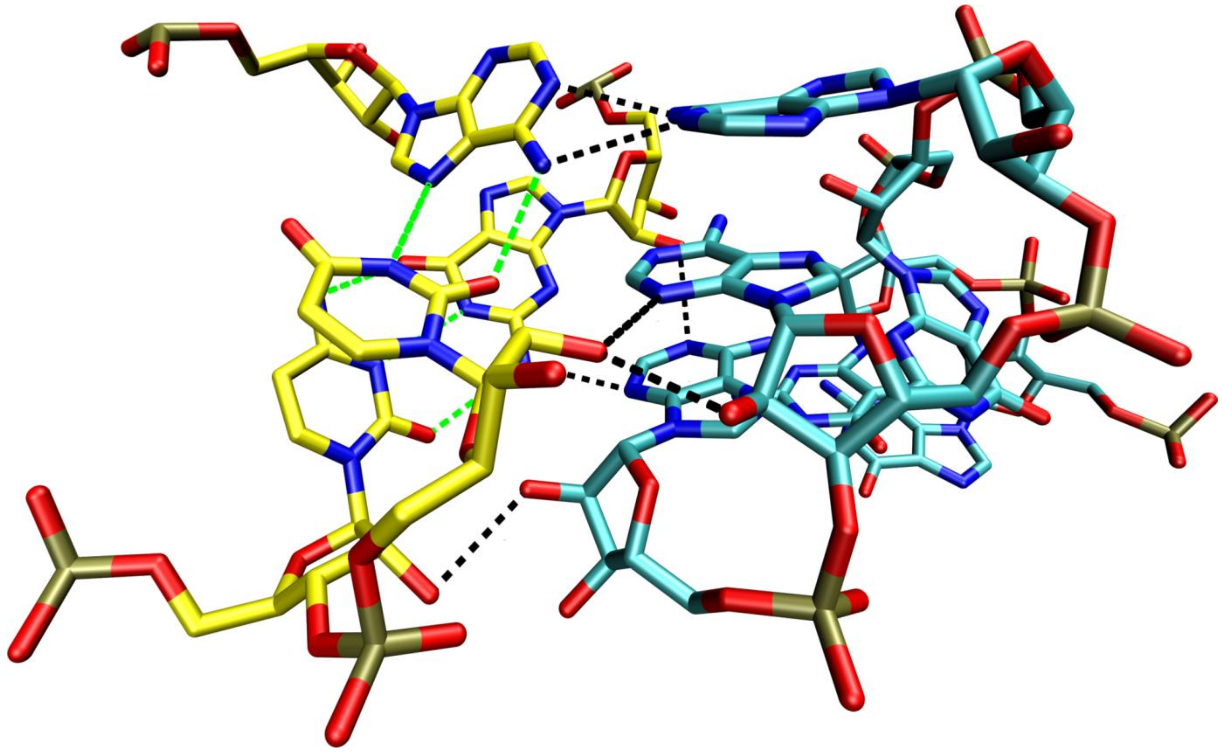

Figure S2. **Definition of the hot-zone region in REST2 enhanced-sampling simulations of the hTTR system.** The hot zone included only the *t*HW A6:U36 and *c*WW G8:C35 base pairs of the tetraloop receptor (carbons in yellow). To maintain their internal stability, these base pairs (base pairing H-bonds indicated by green dashed lines) were stabilized with a 2 kcal/mol sHBfix potential between the hydrogen and acceptor. This setup ensured that enhanced sampling targeted primarily the tertiary interactions with the GAAA tetraloop (carbons in cyan; H-bonds indicated by black dashed lines), without excessively perturbing the internal structures of either the tetraloop or the tetraloop receptor.

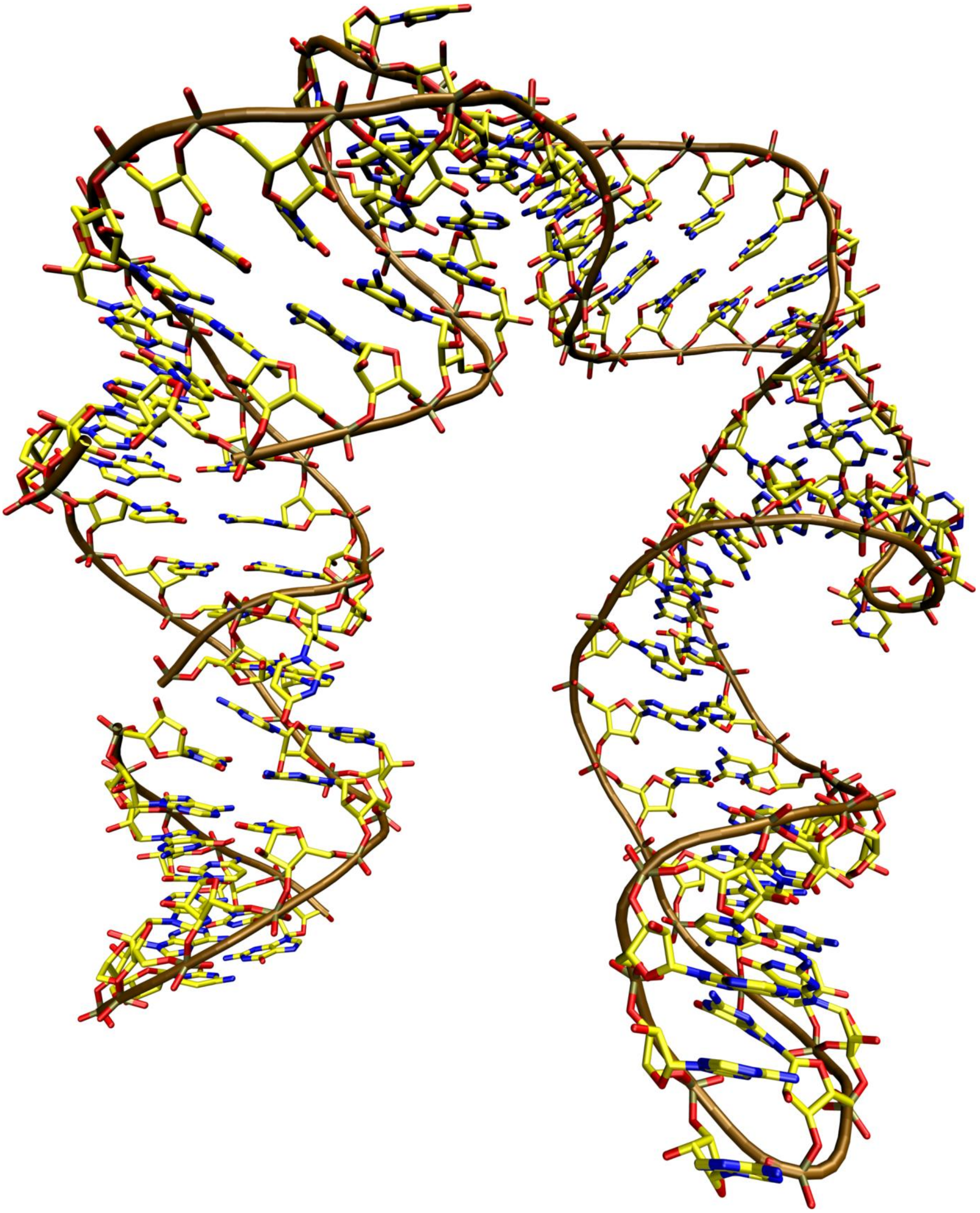

Figure S3. **Structure of the disrupted miniTTR-6 used as start for further MD simulations.** Compare with Figure S1. Shown is a snapshot from an OL3/OPC simulation, later used as the starting structure for additional OL3/SPCE and DES-Amber/TIP4PD simulations. In the OL3/SPCE simulations, the native RNA fold was eventually restored (see main text and Figure S5).

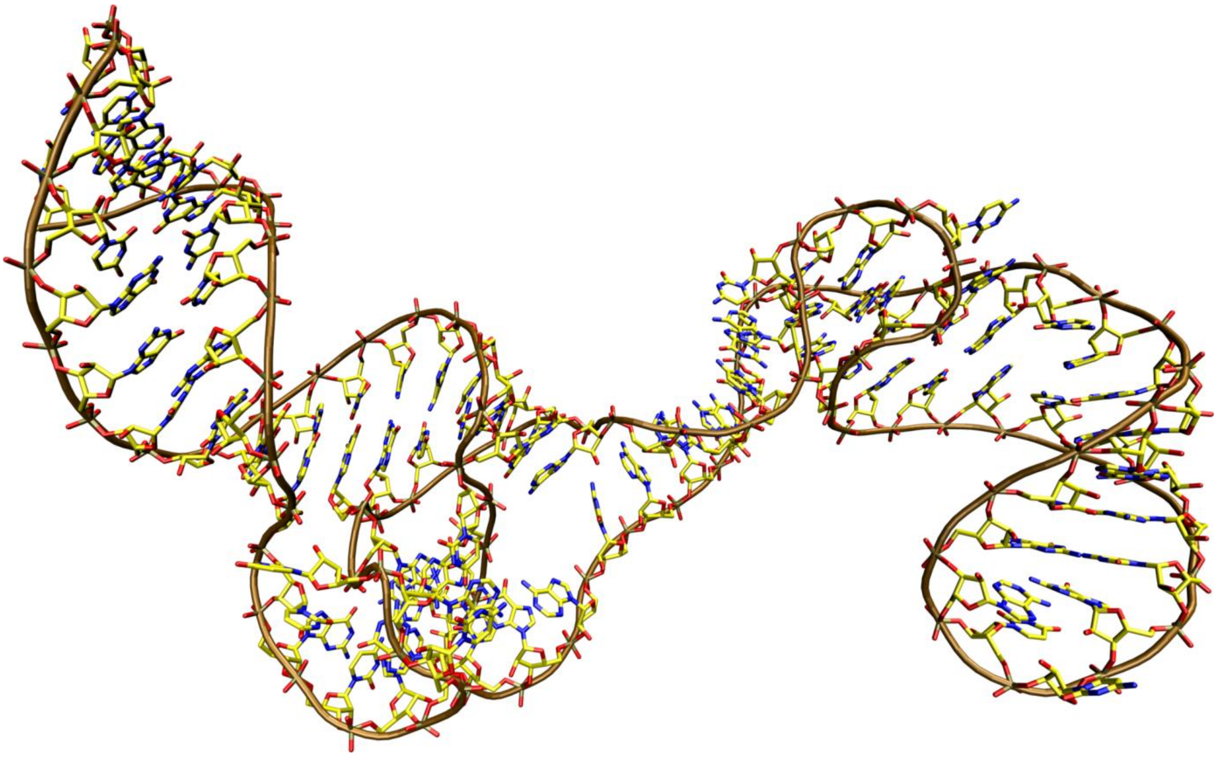

Figure S4. **Structure of the disrupted H.m. L1 stalk rRNA.** Compare with Figure S1. Shown is a snapshot from DES-Amber/TIP4PD simulations, in which the largest distortions were observed for this system. Many native non-canonical rRNA elements tended to become disrupted, giving way to an extended quasi-A-form structure.

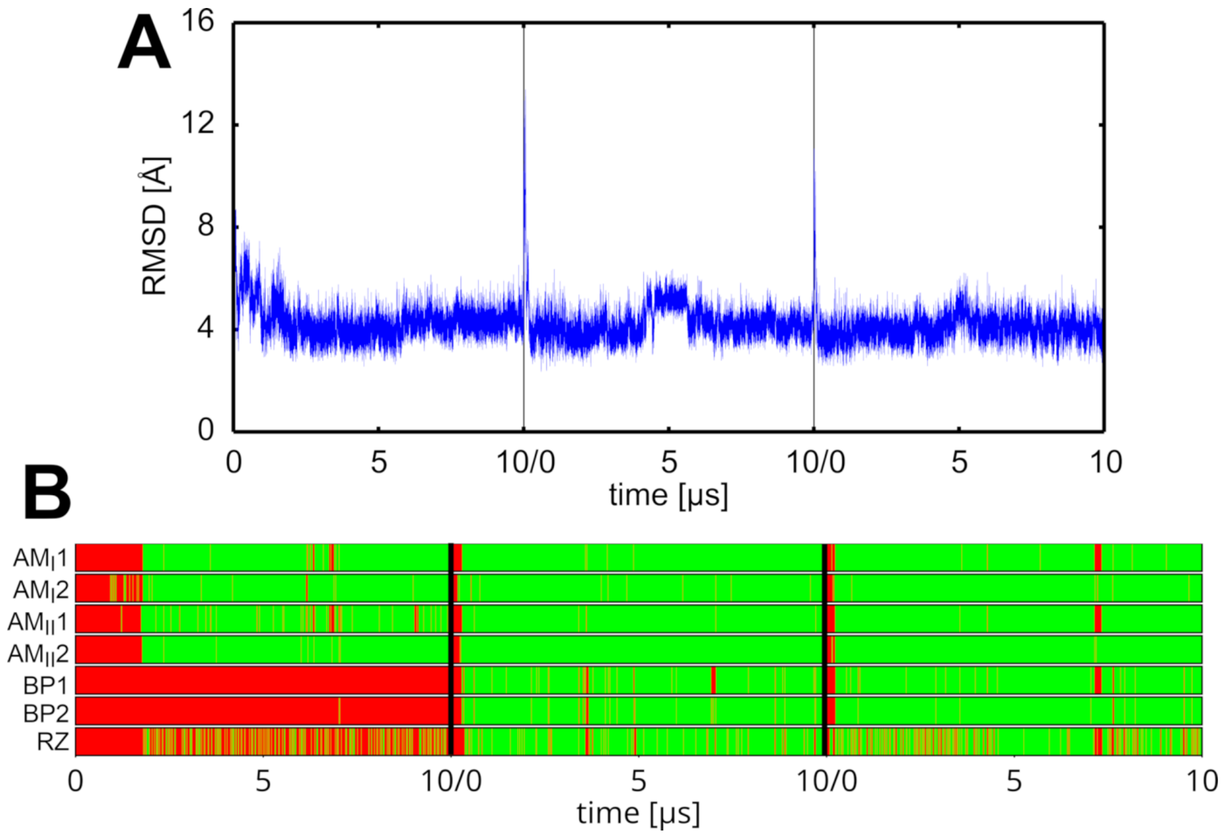


Figure S5. **Restoration of the native fold of miniTTR-6 in additional SPC/E simulations, started from a structure previously disrupted in OL3/OPC simulations. A)** Time evolution of the RNA RMSD in the restoration simulations. **B)** Time evolution of the signature TTR H-bonds. Green and red indicate presence and absence of the H-bond, respectively. A full restoration of the TTR was observed in two of the simulations and a partial restoration in one. See main text Figure 3 for definition of the individual H-bonds. See Figure S3 for the starting structure.

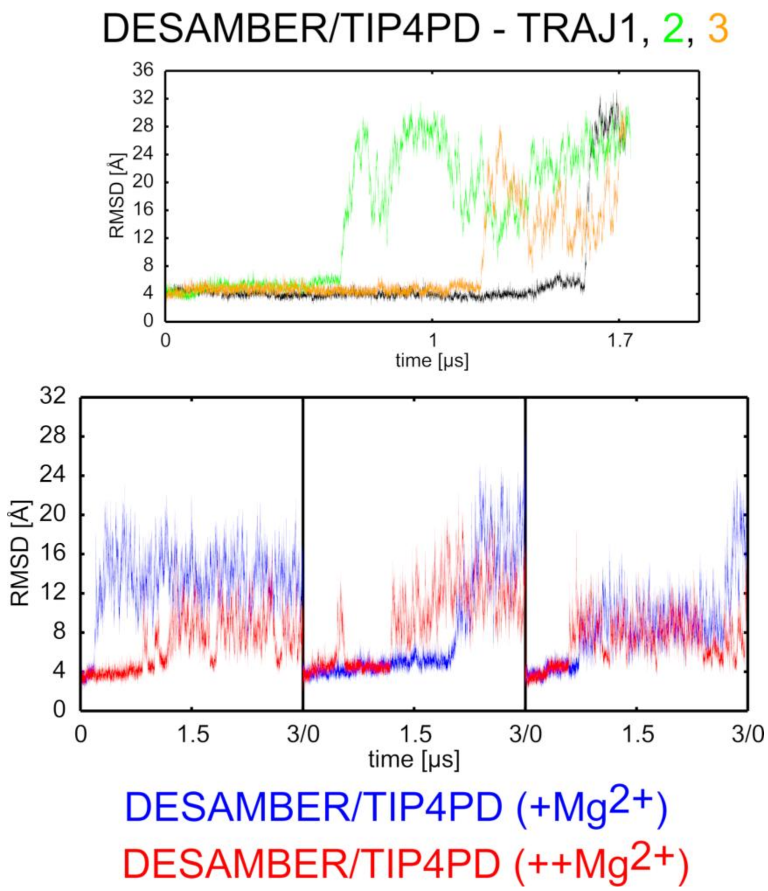


Figure S6. **Loss of the native RNA fold in simulations of miniTTR-6 using the DES-Amber FF and TIP4PD water model.** Time evolution of the RNA RMSD in simulations using different ionic conditions. Datasets are color-coded according to the legend.

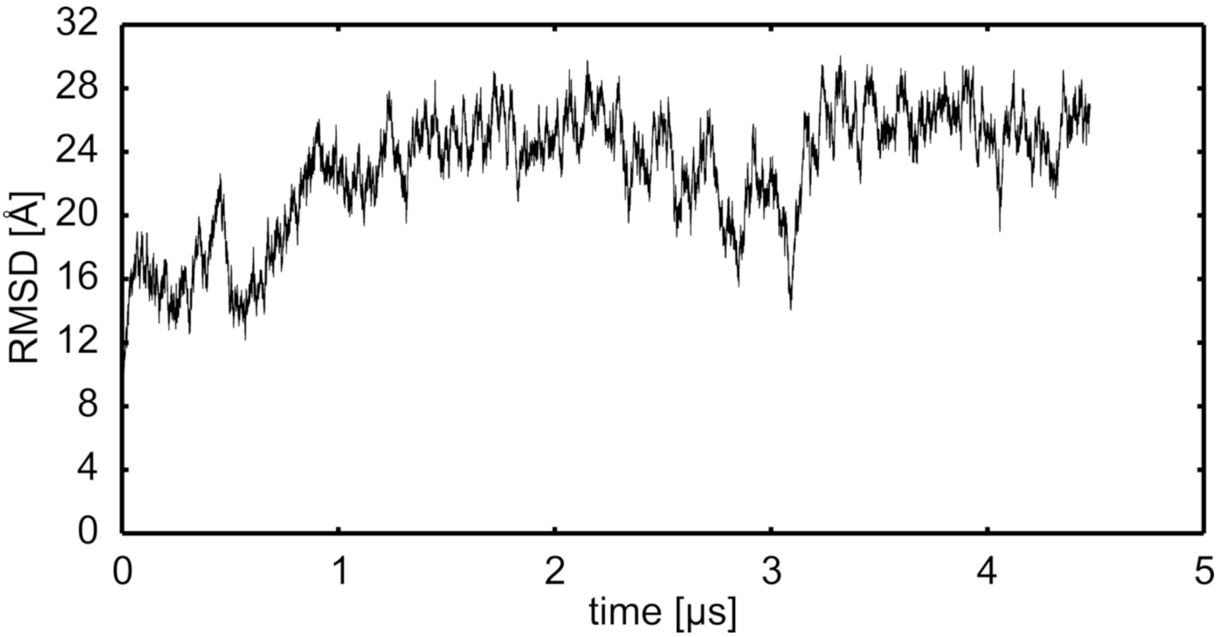


Figure S7. **Unsuccessful attempt to spontaneously restore the native fold of miniTTR-6 in DES-Amber/TIP4PD simulation.** Although the initial box size was substantially increased to prevent image clashes, the system showed no tendency to return to the native fold and instead continued to deviate further from it.

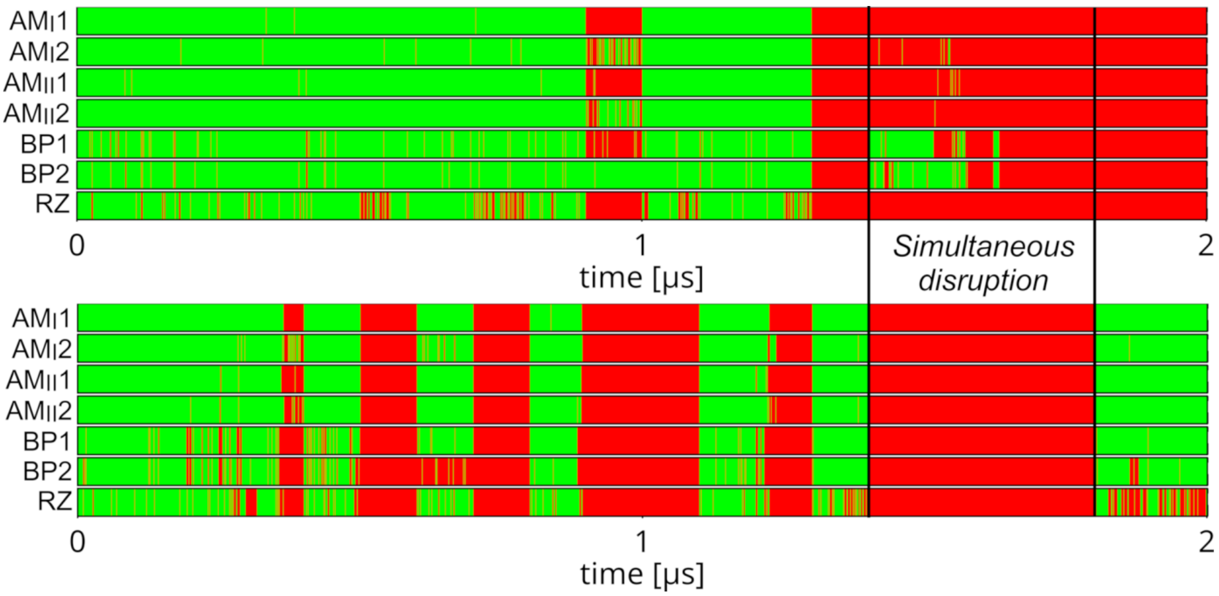


Figure S8. **Simultaneous disruption of the two TTRs in REST2 OL3/OPC simulations of hTTR.** Time evolution of the signature H-bonds of the TTR motifs in a demultiplexed continuous trajectory from the REST2 simulations with the OPC water model. Both TTRs are shown (top and bottom). The black bars and label highlight the instance of simultaneous disruption of both motifs, followed by full spontaneous restoration of one. Green and red indicate presence and absence of individual hydrogen bonds, respectively (see Figure 3 in the main text for definitions).
